# A deep neural network to de-noise single-cell RNA sequencing data

**DOI:** 10.1101/2024.11.20.624552

**Authors:** Mohsen Sharifitabar, Shiva Kazempour, Javad Razavian, Sogand Sajedi, Soroosh Solhjoo, Habil Zare

## Abstract

Single-cell RNA sequencing (scRNA-seq), a powerful technique for investigating the transcriptome of individual cells, enables the discovery of heterogeneous cell populations, rare cell types, and transcriptional dynamics in separate cells. Yet, scRNA-seq data analysis is limited by the problem of measurement dropouts, i.e., genes displaying zero expression levels. We introduce ZiPo, a deep artificial neural network for rate estimation and library size prediction in scRNA-seq data which incorporates adjustable zero inflation in the distribution to capture the dropouts. ZiPo builds upon established concepts, including using deep autoencoders and adopting the Poisson and negative binomial distributions, by taking advantage of novel strategies, including library size prediction and residual connections, to improve the overall performance. A significant innovation of ZiPo is the introduction of a scale-invariant loss term, making the weights sparse and, hence, the model biologically more interpretable. ZiPo quickly handles vast singular and mixed datasets, with the processing time directly proportional to the number of cells. In this paper, we demonstrate the power of ZiPo on three datasets and show its advantages over other current techniques. The code used to produce the results in this manuscript is available at https://bitbucket.org/habilzare/alzheimer/src/master/code/deep/ZiPo/.

## I. Introduction

RECENT single-cell RNA sequencing (scRNA-seq) technologies provide invaluable insight into gene expression at individual cell levels. However, they often result in many genes displaying zero expression levels. These zero values can arise from two distinct events: (i) biological events, referred to as structural zeros [1], when a gene expresses no RNA at the time of the experiment, and (ii) non-biological events, also known as dropout events [1]–[ stemming from the insufficiencies in library preparation and amplification processes [4], [5]. Non-biological zeros occur due to various factors in the employed technology. These factors can include mRNA degradation after cell lysis, variability in converting mRNA to cDNA, inconsistencies in amplification efficiency, cell library concentration or sequencing depth, and other batch effects [3].

Unique Molecular Identifiers (UMIs) are a valuable tool in rectifying amplification biases in non-zero gene expression measurements by identifying and eliminating reads from cDNA duplicates generated during amplification [6]. Despite their utility, UMIs cannot rectify sampling zeros and the corresponding cDNA copy numbers for these zeros remain unidentified [7].

In the context of scRNA-seq data, imputation involves inferring missing expression values or non-biological zeros. The aim is to estimate the actual expression values that were not detected due to experimental limitations. Imputation methods are often developed based on the principle that similar cells will likely have similar gene expression profiles [8], [9].

Noise in scRNA-seq data can arise from various sources, including the aforementioned technical variabilities and the inherent stochasticity of gene expression. Denoising methods strive to eliminate or reduce noise in the data by identifying and adjusting for these sources of variability, making the signal more pronounced. For instance, a common denoising approach adjusts the expression values for each gene based on the total amount of RNA detected in each cell, thereby correcting for differences in sequencing depth [10]. Statistical learning and deep learning models, such as autoencoders, are among the most powerful tools for processing complex biological data like scRNA-seq.

Imputation methods for scRNA-seq data are generally built upon probabilistic and mathematical models. They aim to replace missing values, often zeros, with estimates derived from the existing data points and their interrelationships. To infer the missing values, these methods often exploit the structure and characteristics of the data, such as the correlation patterns in the scRNA-seq data, i.e., the similarities among cells and genes.

One example is the Markov affinity-based graph imputation of cells [11] (MAGIC), which is among the most renowned statistical methods tailored for denoising and imputation of scRNA-seq data. This computational approach denoises the cell count matrix and imputes missing gene expression values. MAGIC constructs a graph where nodes represent cells and edges reflect similarities between cells, facilitating information sharing across similar cells via a diffusion process. Another notable method, SAVER [12], uses an empirical Bayes-like approach to estimate the true expression levels, leveraging expression similarities between cells to impute non-biological zeros. scImpute [8] is another method that finds and imputes highly probable dropout values using a clustering approach. This method draws on information from the same gene in similar cells to the imputed cell. A similar approach DrImpute [13], employs k-means clustering [14], [15] to identify similar cells and then imputes dropouts based on the average expression values in those cells.

Deep learning models have emerged as revolutionary tools in the last two decades due to their unique ability to solve complex problems. With the exponential growth of biological data, these models help determine patterns and draw inferences from large, complex datasets. Deep models leverage artificial neural networks with multiple hidden layers to perform downstream tasks, including feature extraction, dimension reduction, prediction, and classification. They offer novel approaches to tackle and interpret the complexity of this information, providing new insights into various areas such as genomics, proteomics, and cell biology [16]–[18]. Applications vary from predicting protein structure and detecting genetic variants to understanding gene regulatory networks and interpreting scRNA-seq data. As these models evolve, their application in biological research is expected to drive significant breakthroughs and help reveal the complex underlying mechanisms.

Notably, self-supervised models like autoencoders play a prominent role in deep learning. Deep autoencoders are explicitly designed to provide efficient data representations. First, the encoder compresses the input into a lower-dimensional representation. Next, the decoder reconstructs the original data from the compressed representation. Training an autoencoder involves minimizing the difference between the input and the output. Incorporating multiple layers in the encoder and the decoder results in deep autoencoders that can model complex data with a high-dimensional feature space. However, a challenge known as the “vanishing gradients” emerges during the backpropagation stage of training, particularly in deeper models. Because of the chain rule and multiplication of the gradients, there is a possibility that when some of the gradients are small, the multiplication would be numerically too small, leading to no significant learning in back-propagation. Techniques such as batch normalization [19] and residual connections [20] fix the vanishing gradients problem [21]. Deep autoencoders are valuable in different applications, including feature extraction, dimension reduction, and denoising.

Regardless of the denoising approach, discrete statistical distributions are needed to model the count data in scRNA-seq. Various count distributions such as negative binomial (NB), zero-inflated NB (ZINB), Poisson, and zero-inflated Poisson (ZIP) have been widely employed for modeling UMIs or gene reads in scRNA-seq [4]. For instance, the deep count autoencoder (DCA) [22], is used to denoise the scRNA-seq data using an NB model for noise. This approach characterizes the reconstruction error by evaluating the probability distribution of the noise model rather than directly reconstructing the input data. The ZINB distribution introduces a zero-inflation parameter that signifies the proportion of additional zeros, distinct from those originating from the NB component. Similarly, the ZIP distribution also incorporates a zero-inflation parameter, specifically targeted at the zeros not accounted for by the Poisson distribution. Employing a zero-inflated model versus a non-zero-inflated model does not depend on the prevalence of zeros in the data. Zero inflation is a statistical consideration that takes into account the inevitable nonbiological zeros. Zero inflation and its biological and technical validity for different sequencing methods are among the highly debated topics.

Here, we introduce ZiPo, a deep model that takes scRNA-seq data as its input and performs two tasks: (a) imputing the missing data using zero-inflated distributions, and (b) providing an embedding of the single-cell gene expression data in a low-dimensional space, offering a nonlinear transformation akin to PCA’s dimensionality reduction capabilities that can be used in downstream tasks like cell-type clustering. We introduce a regularization parameter that not only adjusts the zero-inflation level but also offers a more refined, adaptable, and biologically accurate approach for handling the intricacies of sequencing data, setting our work apart in this domain. This model can also be used for single-nucleus data, but we only use single-cell terminology in this manuscript for simplicity.

## II. Methods

In ZiPo, we first encode the high-dimensional gene expression data of the cells into a low-dimensional space (e.g., with 32 latent variables) using a deep model. Next, we consider these latent variables as unknown covariates of the scRNA-seq data and recover the expression levels using these unknown covariates together with other possibly known covariates that may influence the gene expression patterns observed in individual cells. Such covariates might include tissue or cell types, sequencing depth, spatial location, age, sex, etc. [23].

We assume a zero-inflated distribution for gene expression levels. The two most common distributions for scRNA-seq data are *zero-inflated Poisson* (ZIP) and *zero-inflated negative binomial* (ZINB). The results in this manuscript are based on ZIP, but they can readily apply to ZINB with minimal adjustments. We prioritized ZIP because the straightforward Poisson distribution aligns well with UMI-based sequencing data. This approach eliminates the need for ZINB’s additional parameter that might account for potentially greater dispersion in the model.

The model takes each cell’s measured gene expression vector as its input. The encoder consists of dense linear layers for reducing the dimension followed by sigmoid activation. The model is then followed by parallel layers from the encoded variables (unknown covariates) to predict:

1. the library size,
2. the zero-inflation rate, and
3. the parameters of the desired distribution.

A schematic representation of the model is provided in Fig. 1.

**Figure 1:**
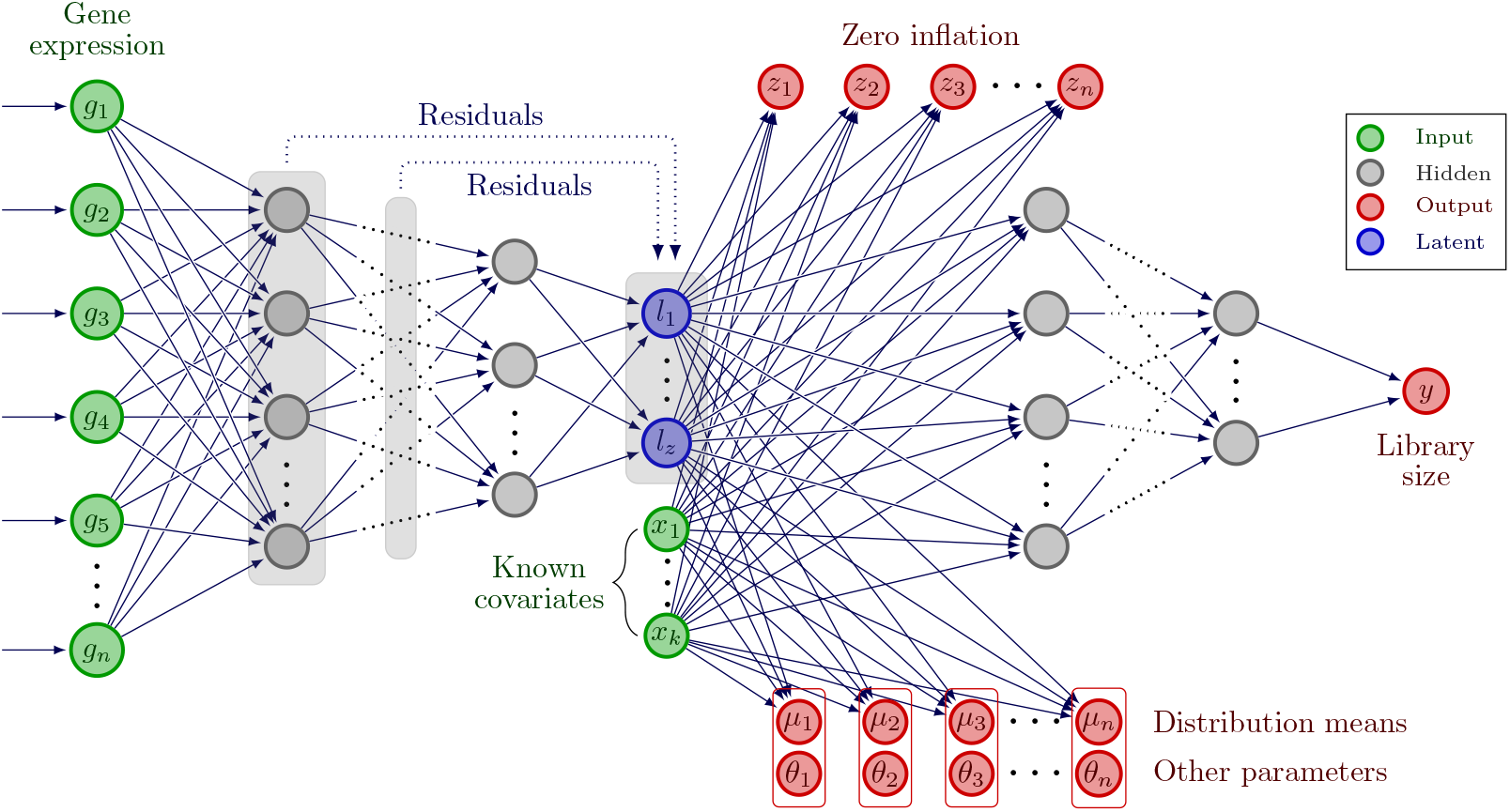
Schematic of the ZiPo model structure. ZiPo is composed of a deep encoder and a shallow decoder. Each input sample is a vector of expression of *n* genes *g*_1_,, *g*_*n*_ for a given cell. Residuals can optionally be added from any encoder layer to the layer containing latent variables *l*_1_, …, *l*_*z*_. If there are any known covariates *x*_1_,, *x*_*k*_, we feed them to the model alongside the latents. The decoder network has several hidden blocks, each composed of standard components including dense linear, activation, and batch normalization layers. The decoder generates zero-inflation probabilities *z*_1_, …, *z*_*n*_, the distribution means *µ*_1_,, *µ*_*n*_, and other possible distribution parameters *θ*_1_,, *θ*_*n*_. Also, a deep subnetwork will be used to predict the library size, *y*, using the latents and covariates.

### A. Advantages of predicting the library size

We separate the library size (i.e., the sum of all gene expressions for each cell) from the other predictions because what we want to model in the zero-inflated distribution is the relative expression levels of genes in a cell, not the absolute values, which depend on less relevant variables like the sequencing quality, assay efficiency, etc. To normalize gene expression data, we can adjust raw count data using count per million (CPM) [24] to account for differences in library size or sequencing depth between samples, allowing for fair comparisons of gene expression levels across samples. In this method, the raw counts for each gene in a sample are divided by the total number of reads (or counts) in that sample. Predicting the library size offers at least two distinct advantages over calculating it via adding all gene expressions for each cell: (1) it can help reduce technical noise in the data, and (2) it retains potentially valuable information about the library size in the latent variables that can be used in subsequent analyses. Notably, the library size could convey information relevant to the cell size [25].

### B. Sparsifying of the weights

We expect each feature in the second layer to represent a pathway or an eigengene, incorporating only a small subset of the genes [26], [27]. Therefore, most of the weights of *the first encoding layer* should be zero or near zero, forming a sparse weight matrix. The sparsity of this layer can be enforced by adding a term to the loss function that acts similarly to an *L*^1^ penalty. Not all activation functions can produce the intended sparsity as discussed in detail in section II-D.

### C. The model structure in mathematical notation

Throughout the manuscript, we use the subscript *c* for any variable to indicate that it is calculated for the cell *c*. The input matrix is denoted as *X* = {*x*_*cg*_}, where *x*_*cg*_ is the expression of gene *g* in cell *c. X*_*c*_ denotes the full expression vector of cell *c* over all available genes. For *t* encoding layers, the cellular embedding for cell *c* is formulated as:

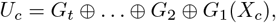

where *U*_*c*_ is the embedding for cell *c*. Each *G*_*i*_ is composed of a linear layer, an activation function (e.g., sigmoid), and a *batch normalization* layer, which helps resolve the vanishing gradients problem and accelerates the training [19], [28]. We add known covariates of the cells, *U*_*c*_^*′*^;, to the embedding vector *U*_*c*_, yielding the complete encoded representation of the cell,

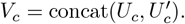

Subsequently, optional layers are used to predict the following variables:

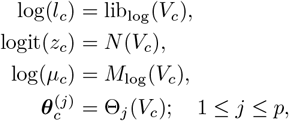

where *l* is the library size, *µ* is the mean parameter for the distribution, *z* is the zero-inflation probability, ***θ*** represents additional relevant parameters of the distribution, and *p* is the number of these parameters. In the case of the Poisson distribution, we only have the mean parameter, *µ*. For the NB distribution, we need an extra parameter representing the difference between the mean and the dispersion of gene expressions. For *N*, we use a dense linear layer. For *M*_log_, we use a dense linear layer followed by *logSoftmax* activation, forcing the sum of the average gene expressions for each cell to be equal to 1.

Given this configuration, the following linear relationships are established:

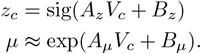

To model the library size, we use a series of dense linear layers followed by batch normalization and an activation function. We can readily employ deep models to predict distribution parameters, resulting in nonlinear rather than linear *V*_*c*_ functions in the above equations.

### D. The loss function

The zero-inflated distribution is given by:

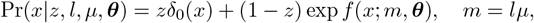

where *f* represents the logarithm of the distribution for non-zero-inflated expression levels with mean *m* and possibly other parameters ***θ*** and δ_0_ is the atomic distribution on zero. Note that *m* = *lµ* is the expected expression level according to our definitions.

We denote the rate of zero inflation by *r*, i.e., *r* = logit(*z*) and the likelihood of our samples, given the distribution parameters by ℒ. The first component of the loss function is the log-likelihood of the zero-inflated distribution:

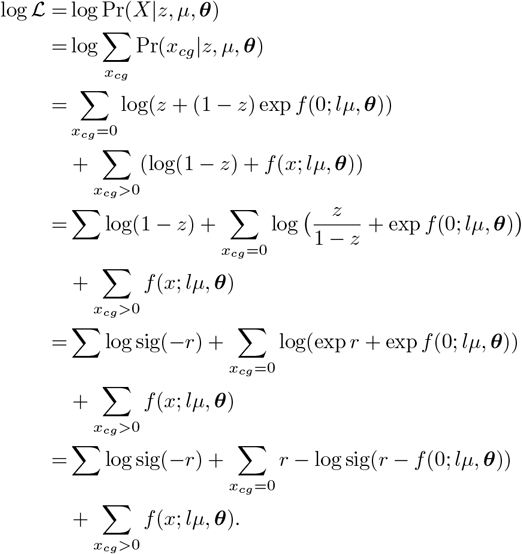

A typical approach employed in the DCA method involves using truncated log functions by adding a tiny offset quantity to the argument of the log function to avoid infinity (or very large) numeric values when the argument is zero (or near zero). Although this approach might be appropriate in specific conditions, it is inaccurate when we have no a priori knowledge of the input argument to decide on a proper value for the offset. To prioritize precision in our calculations, instead of relying on truncated log functions, we use the composite function log sig [29], which stabilizes the numerical schema without truncation.

The ZiPo model is regularized in two ways. We apply *L*^2^ regularization to the predicted zero probabilities and other distribution parameters. The regularization hyperparameter in ZiPo, *α*_*z*_, controls the level of zero inflation. An excessively large *α*_*z*_ prevents the model from using significant zero inflation. Setting *α*_*z*_ to zero will yield a model with strong zero inflation that overestimates the expression rates. We also impose a regularization term on the weights of the first layer to sparsify them. We use this regularization term to shrink weights appearing to be less important. We then prune those relatively small weights after ensuring that removing them does not affect the model performance.

We use NLL for “negative log-likelihood” and MSE for “mean square error” throughout the manuscript.

### E. Which activation function and weight regularization approach should be used?

In deep neural networks, activation functions such as *ReLU* [30] or *GELU* [31] are commonly employed to tackle the vanishing gradients issue. However, for the encoding segment of ZiPo, we favor the sigmoid activation function. This choice is influenced by the specific type of regularization applied to the weights (parameters) of the first encoding layer.

A limitation of ReLU arises from its linear properties. That is, the model can decrease the absolute weights in the initial layer while compensating for this by increasing the weights of the second layer, without affecting the final output of the second layer and the final performance of the model. Therefore, simple *L*^1^ or *L*^2^ regularization penalties would not significantly change the model’s performance or output. Some variations of the above activation functions like *ReLU6*, defined as min(max(0, *x*), 6), could potentially resolve this problem. However, we prefer the nonlinearity of the sigmoid function over the piecewise linearity of *ReLU6*.

Similarly, the optimizer can lower the weights of *the first encoding layer* but compensate for them by increasing the batch normalization weights in the following layer. To protect the first layer weights, we use a scale-invariant loss function defined by dividing *L*^1^ by *L*^2^:

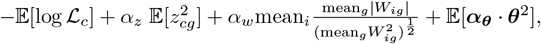

where *W*_*ig*_s are the weights of the first linear layer and *α*’s are regularization hyperparameters determining the strength of the corresponding penalty terms (i.e., *α*_*z*_ controls the zero-inflation rate and *α*_*w*_ controls the weights in the first layer and *α*_***θ***_ controls the parameters of the distribution).

If the features of the initial layer are sufficiently interpretable, the model should perform well because those features are directly related to the genes. Employing similar regularization across all layers is possible. However, introducing an additional hyperparameter for each layer complicates the model and the training process.

### F. Datasets

We have used three datasets to illustrate ZiPo performance (Table I). The ROSMAP Alzheimer’s disease single-nucleus RNA-seq (snRNA-seq) dataset was generated by Mathys et al. [32], and is available from https://www.synapse.org/ with synapse ID: syn18485175. This dataset includes 80,660 single nuclei derived from the dorsolateral prefrontal cortex of 48 postmortem human brains. The liver [33] and myeloma [34] scRNA-seq datasets are accessible through GEO with accession numbers GSE125188 and GSE117156, respectively. In the liver dataset, 70,706 human CD45+ cells from the paired liver perfusion, spleen, and human peripheral blood mononuclear cells (PBMC) were profiled. The myeloma dataset includes 51,840 cells from 29 diagnosed patients and 11 control donors.

**Table I:** Three datasets were used in our experiments testing the ZiPo model.

| Dataset | Tissue(s) | # Cells | # Genes | Nonzero |
| --- | --- | --- | --- | --- |
| ROSMAP | Brain | 70,634 | 17,249 | 9.82% |
| Liver | Liver, Spleen, Blood | 68,100 | 19,536 | 9.00% |
| Myeloma | Bone marrow | 51,840 | 27,992 | 1.94% |

While the distributions of library sizes (i.e., the sum of gene expression per cell) were similar across these three datasets, the total gene expression (i.e., the sum of expression levels per gene) varied (Fig. 2). We used the liver dataset to conduct hyperparameter analysis. Then, we assessed the performance of the optimized architecture in all three datasets.

**Figure 2:**
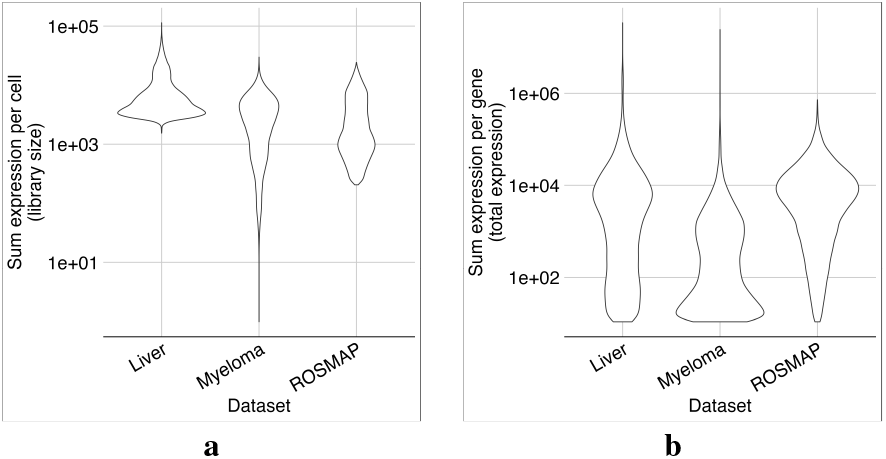
Violin plots of library sizes and gene expression levels for each dataset. Library sizes generally vary by two orders of magnitude. The Myeloma dataset has outliers with relatively small library sizes **(a)**. While the total expression levels of most genes are in the order of 10^4^ in the Liver and ROSMAP datasets, it has a bimodal distribution in the Myeloma dataset **(b)**.

### G. Model parameters

Here, we systematically investigate the hyperparameter space to identify reasonable values and provide insight for ranges that can be extensively explored using an automatic hyperparameter optimization software such as *optuna* package [35]. Specifically, we use two middle layers with 16 and 4 neurons to predict the library size. The *Adam* optimizer [36] is used with the default linear layer initialization of weights in *pytorch* [37]. The desired learning rate is 0.001. However, due to the relatively large library sizes, the optimizer needs large learning rates to converge quickly. Therefore, to accelerate the training process, we set the initial learning rate at 0.0212 and decrease it by a factor of 0.9 to reach 0.001 at epoch 30. If the learning process halts, i.e., no improvement of at least 0.1% in validation loss, the learning rate will be reduced automatically after 15 epochs using the class ReduceLROnPlateau of *pytorch-ignite* [38]. The learning process stops early if the validation loss does not improve over 30 epochs.

In our implementation, users can either manually specify the model hyperparameters or allow our program to search for the optimum settings in a predefined sampling space using the *Optuna* package.

### H. Model comparison

We compared the trained models based on the average loss they obtained using 12 random seeds. The error bars in the figures in the Results section represent *λ* times of standard error of the mean, where 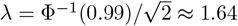, where Φ is the cumulative distribution function for the normal distribution. With this choice, assuming a one-sided *t* test (i.e., normality of distributions and same variance), non-overlapping bars indicate that we can reject the null hypothesis with 99% confidence.

## III. Results

Our systematic investigation of the hyperparameter space starts with seeking an optimum design for the encoder and decoder, using the ROSMAP dataset.

### A. A deep enough encoder with residuals leads to better performance

Almost half of the model variables are embedded in the first layer of the encoder. The first layer also influences the size of the other layers. Therefore, it is important to optimize the first layer. To focus on the encoder, we used a simple linear structure for the decoder, as discussed in II-C.

1. *The first layer’s size:* In a 3-layer model with 64 latent variables, we assessed the effect of the first layer size, *s*_1_, on model training time and performance (Fig. 3). The middle layer size was set to be the geometric mean of the first layer size and the number of the latent variables (i.e., 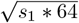). In Fig. 3 and throughout this paper, we use the *s*_1_–*s*_2_– …– *s*_*p*_ convention to describe the number of neurons used in the structure of a model with *p* layers, e.g., 8192–512–64 is a model with 8192, 512, and 64 neurons in the first, second, and third layers, respectively.

**Figure 3:**
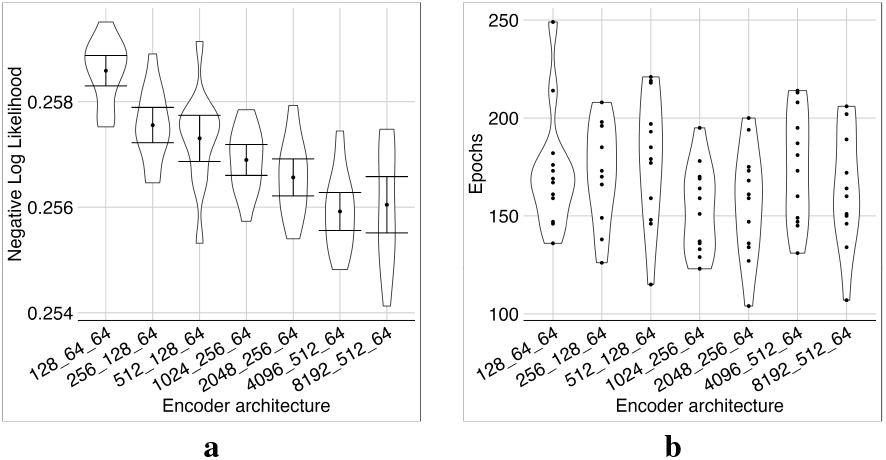
Investigating different sizes for the first layer. We trained each model architecture using 12 different seeds. As expected, the NLL mostly decreased as *s*_1_, the size of the first layer in the encoder, increased **(a)**. The number of training epochs before stopping was roughly in the same range for all encoder architectures **(b)**. We stopped at *s*_1_ = 8192, where the average loss did not improve. *s*_1_ = 4096 had a significantly lower loss than models with a smaller first layer based on the error bars, except for the models with *s*_1_ = 2048. With increasing *s*_1_, the average loss was reduced until *s*_1_ = 4096. The model became relatively unstable, and loss obtained a wider range with *s*_1_ = 8096 (Fig. 3a). The *s*_1_ = 4096 model had the lowest average loss, and according to error bars, was significantly better than the models with a smaller first layer, except the models with *s*_1_ = 2048. Because we are interested in smaller networks, for the rest of our experiments, we chose *s*_1_ = 2048, a smaller model than *s*_1_ = 4096, but not significantly worse in terms of the likelihood. The rationale was that smaller models require less memory and computation for training and have a better chance of interpretability.
2. *The number of latent variables:* In a 3-layer architecture with a first layer of 2048 neurons and the size of the middle layer determined by the geometric mean as described, the likelihood improved as we increased the number of latent variables on the third layer up to 128 (Fig. 4a). Subsequent increases in the number of latent variables caused no significant improvement. As a smaller quantity of latent variables is computationally beneficial (Fig. 4b), we chose 128 latent variables for our model.

**Figure 4:**
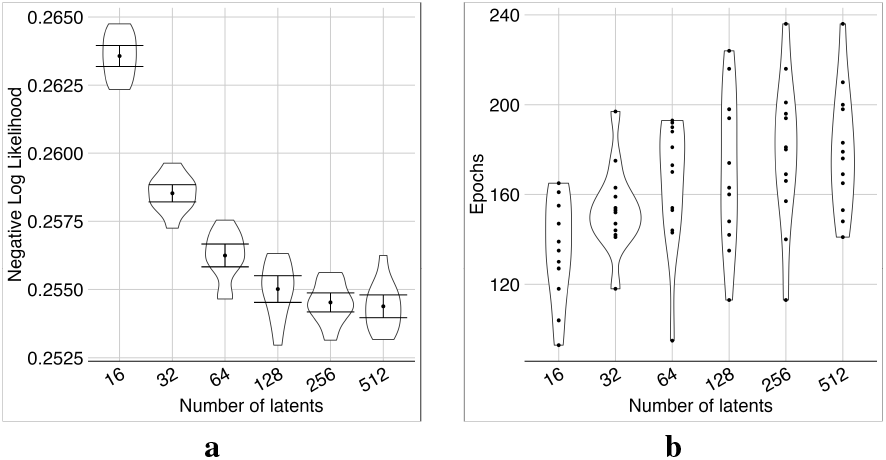
Investigating different numbers of latent variables. Performance improved as the number of latents increased **(a)**. The number of epochs increased slightly as the number of latents increased **(b)**. We stopped the experiment at 512 latents, where the average loss was not improving significantly. The model with 512 latents, had the lowest minimum and was significantly better than the models with fewer latents according to error bars, except for models with 128 and 256 latents. Hence, we chose 128 as the optimal number of latents, which provided adequate performance yet maintained lower complexity compared to larger models.
3. *Deeper encoders with residuals perform better:* We investigated the effect of the encoder depth and the incorporation of residual connections from the encoder layers to the latent layer on models featuring 2048 neurons in the first layer.

Adding residual connections to the models with 3 or more layers lowered their NLL loss (Fig. 5). The deepest model with the full residual connections was the optimal model. Because adding the middle layers did not significantly affect the model size (i.e., most weights are in the first layer), we continued to use this configuration for the rest of our experiments.

**Figure 5:**
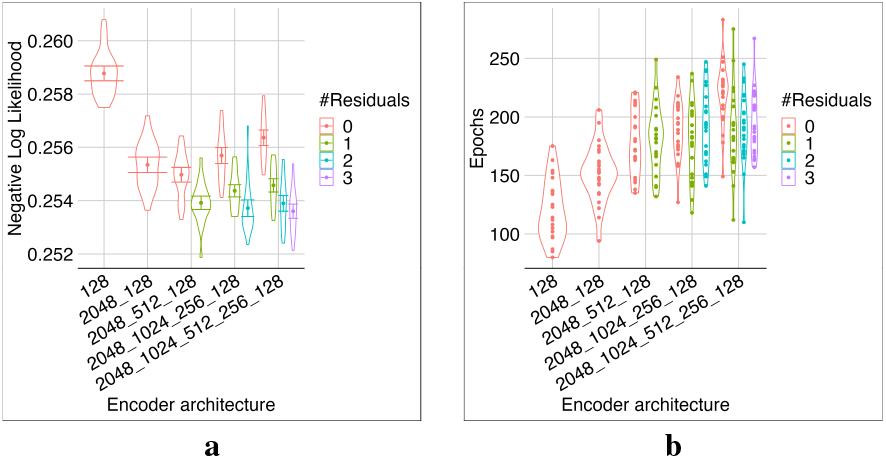
Investigating different encoder depths and residuals. Increasing the model depth improved its performance, but this advantage was lost without residuals **(a)**. The number of epochs required for training was unaffected by the number of residuals **(b)**. Overall, the deepest model with full residuals exhibited superior performance.

### B. A simple decoder is better than a deep one

Using the same optimal encoder structure, (i.e., 2048 neurons in the first layer, 128 latents, 1024, 512, and 256 neurons in the middle layers, respectively, equipped with full residual connections from all encoder layers to the latent layer), we trained the model with different decoder structures (Fig. 6).

**Figure 6:**
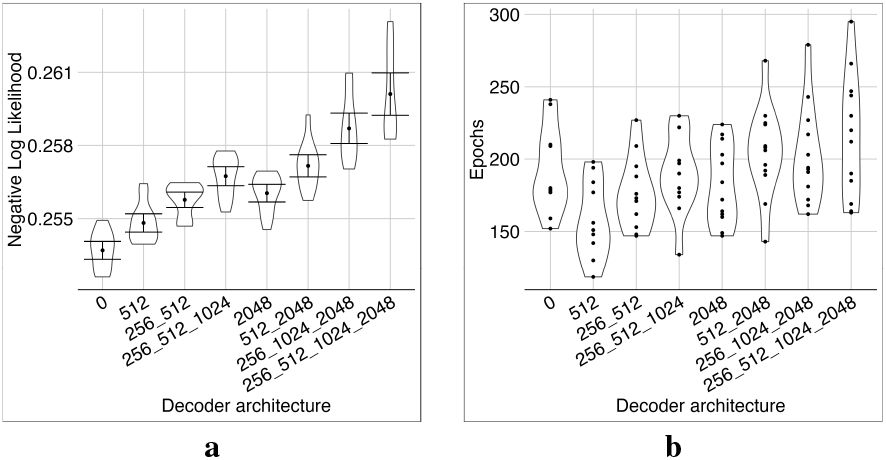
Investigating different decoder architectures. Adding hidden layers to the decoder worsened the model performance **(a)**. Different structures required roughly the same number of epochs for training **(b)**.

More complex decoders needed more epochs for training and resulted in larger NLL losses. Therefore, we chose the simple linear dense decoder described in the Methods section.

### C. Batch normalization is essential for convergence

The effect of batch normalization on training depended on the activation function. With the GELU activation function, removing the batch-normalization layer destabilized the training process, and the optimizer did not converge. With the sigmoid activation function, removing the batch-normalization layer did not destabilize the training process, and the optimizer converged, but got stuck in local minima (Fig. 7). Either way, batch normalization was essential for training.

**Figure 7:**
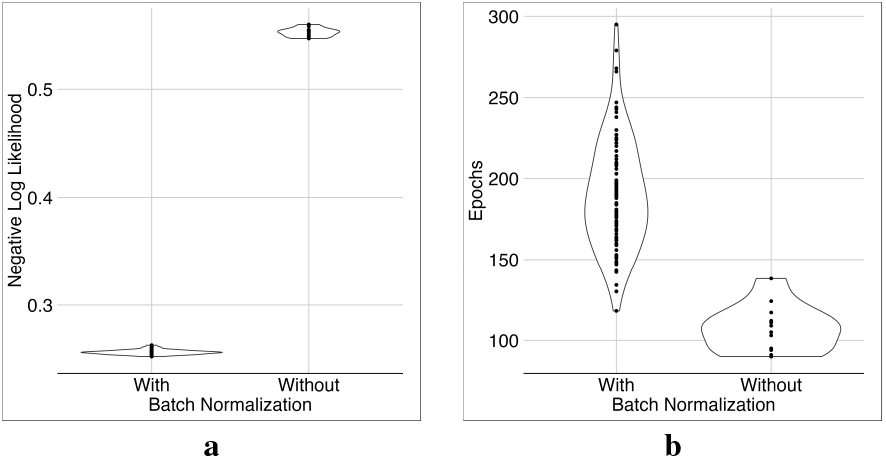
Investigating the effect of batch normalization. Without batch normalization, the model would be stuck in local minima **(a)** and training would stop too early **(b)**.

### D. Controlling the zero-inflation rate improves library size estimation and MSE of reconstruction

With no regularization on zero inflation, the model over-estimated their rates, leading to a much larger MSE in 3 out of 8 tests with different seeds (Fig. 8c). Also, the mean of zero probabilities was very large (Fig. 8d). With a very large zero-inflation regularization parameter (*α*_*z*_ = 10^8^ in Fig. 8d), the model had almost no zero inflation, the training needed more epochs, and the NLL loss was larger than the models with smaller regularization. Apart from these extreme cases, the models had stable reconstruction regarding NLL loss and MSE rates for a vast range of regularization parameters (i.e., 0.01 ≤*α*_*z*_ ≤10, 000). However, smaller regularization parameters generally led to smaller NLL losses and larger mean zero probabilities. In the following experiments, we set *α*_*z*_ = 0.1, which appeared to be a reasonable value for the mean of zero-inflation probabilities.

**Figure 8:**
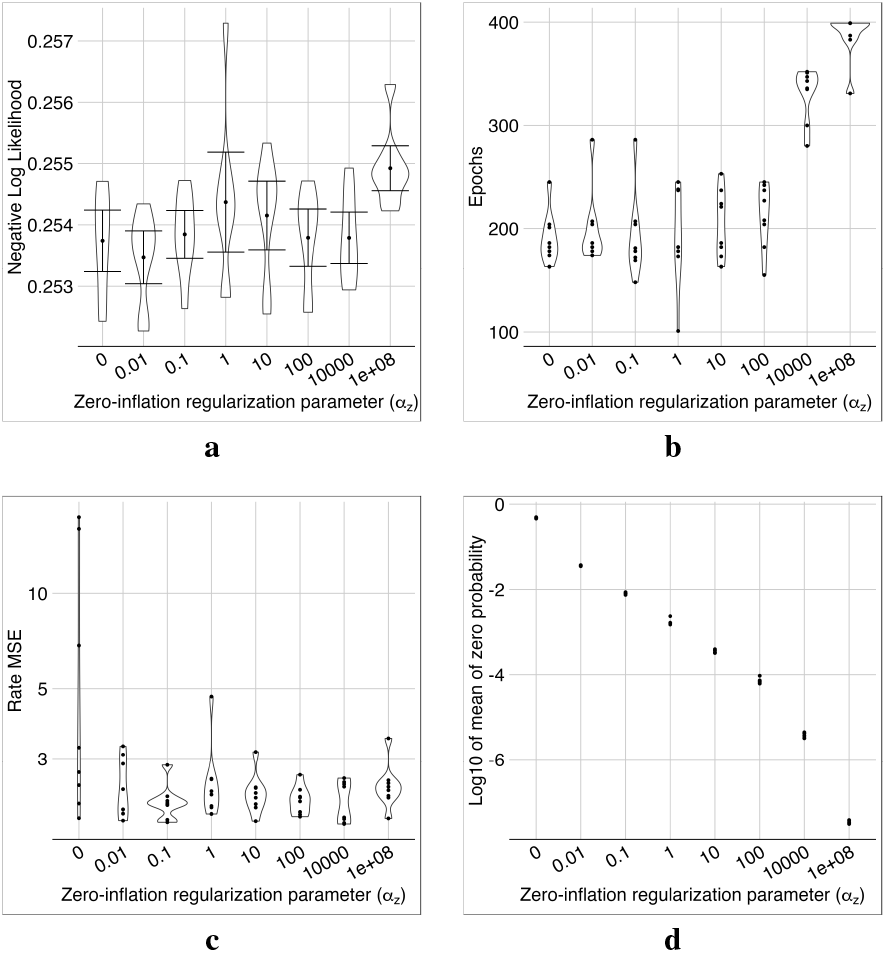
The effect of zero-inflation regularization. NLLs were roughly in the same range except for the very large regularization parameter of 10^8^ **(a)**. The same was true for the number of epochs up to 100 **(b)**. MSE of expression rates was only unstable if no regularization was applied **(c)**. As expected, the mean of zero-inflated probability decreased as the regularization parameter *α*_*z*_ increased **(d)**.

### E. Sparsifying the weights through weight regularization

With *α*_*z*_ = 0.1, we investigated the effect of sparsifying the weights in the first layer using the weight regularization parameter *α*_*w*_. Note that the relative weights are important for each neuron on the first layer. One can multiply all the weights of a neuron by a constant and compensate for it by changing the batch normalization coefficient.

We normalized the weights of each neuron in the first layer by dividing all weights by the maximum weight. Then, we prune the first layer weights by setting those smaller than a threshold δ to zero. We investigated the effect of this sparsification on the model performance.

With no regularization (i.e., *α*_*w*_ = 0), more than 90% of the weights were above 0.01 (Fig. 10a). With a large regularization parameter, say *α*_*w≥*_ 1, many weights were very small, making the model sensitive to pruning (Fig. 10, c and d). In the middle range (i.e., 0.01 ≤*α*_*w*_ ≤0.1), the sensitivity of MSE of rates to the pruning seemed reasonable (Fig. 10c), we had even better NLL losses compared to no regularization (Fig. 9a), and the weight loss stayed almost the same (Fig. 9d).

**Figure 9:**
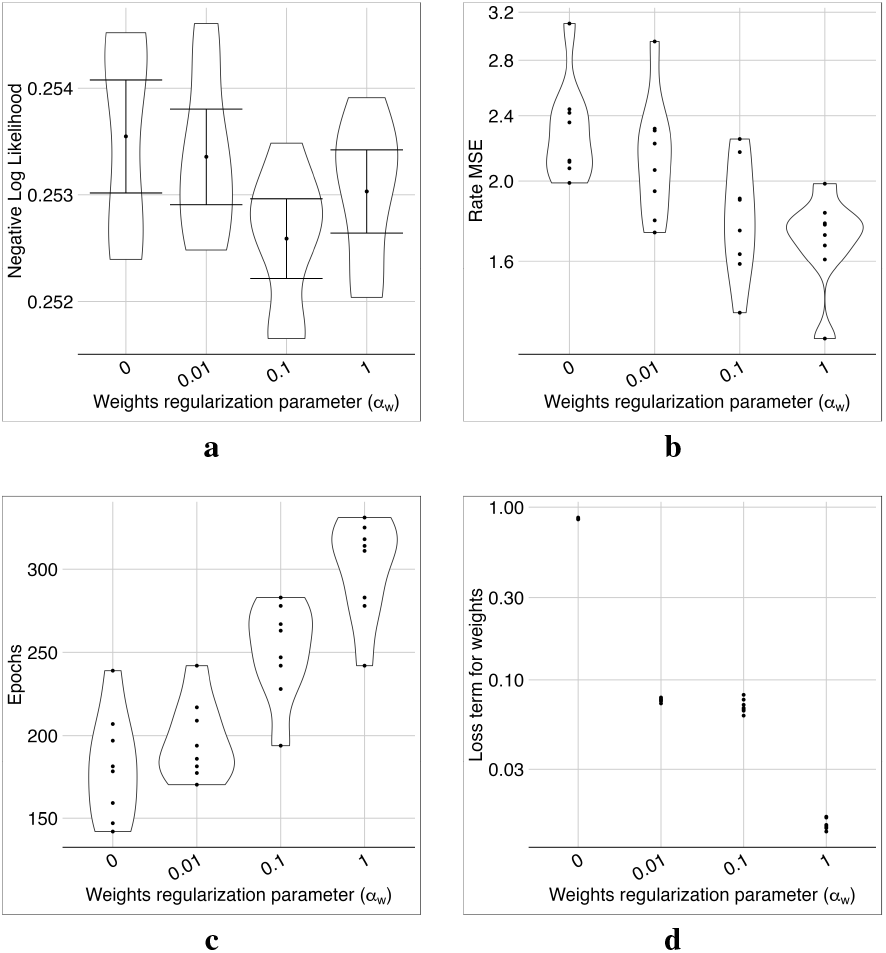
The effects of weight regularization on model performance. Negative log likelihood **(a)** and MSE **(b)** were better in the mid-range of the weight regularization parameter *α*_*w*_ = 0.01 ™1. Training needed more epochs to converge for larger *α*_*w*_ values **(c)**. The corresponding loss term appeared to decrease with increasing *α*_*w*_ and was almost flat in the 0.01–0.1 range **(d)**. Overall, these metrics suggested *α*_*w*_ = 0.1 as an appropriate choice.

**Figure 10:**
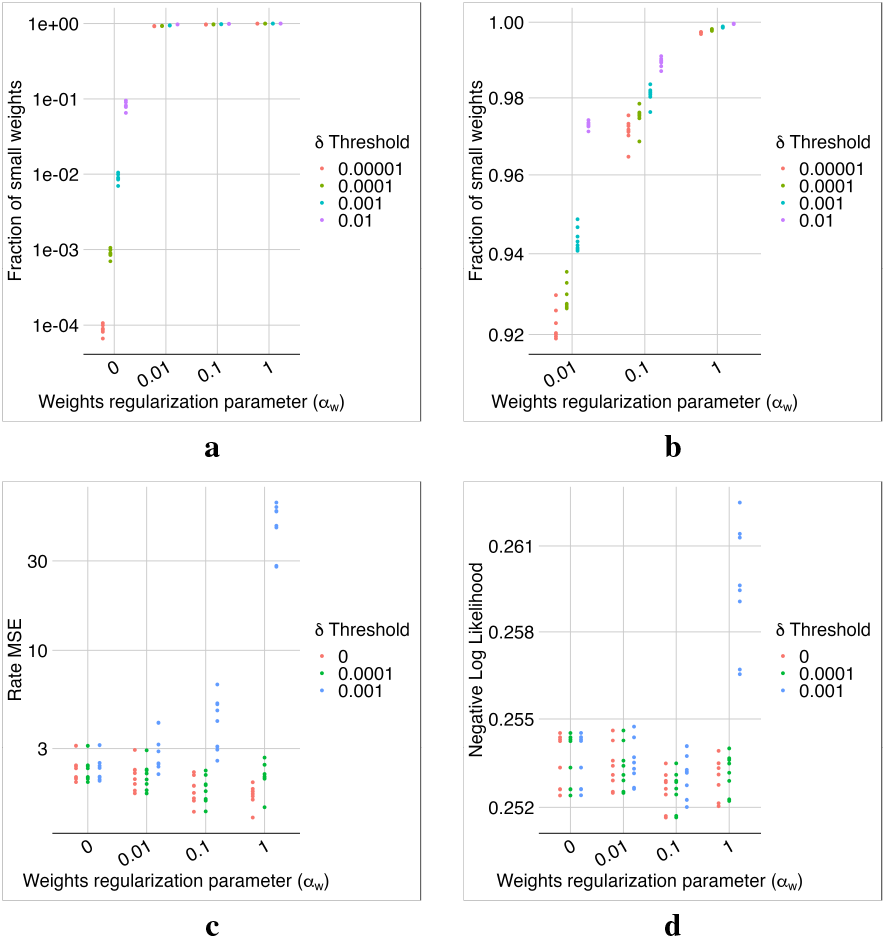
The effects of weight regularization on model sparsity. With no weight regularization, we do not have enough small weights to sparsify the first layer **(a)**. With adding even a small regularization parameter of *α*_*w*_ = 0.01, at least 92% of weights deopped below 10^−^5 **(b)**. The MSE **(c)** and NLL **(d)** losses were more sensitive to the pruning of weights for stronger regularization. Overall, *α*_*w*_ = 0.1 appeared to be a reasonable choice, leading to nearly 98% of the weights becoming zero after pruning with δ = 0.0001 without affecting NLL and MSE losses.

### F. ZINB compared to ZIP

Our experiments showed that due to the greater complexity of the NB distribution compared to the Poisson distribution, we would need to apply another regularization (although very small) on the predicted rates of the model to train it. Therefore, we added 0.1∗ 𝔼 [(*l*_*c*_*µ*_*c*_ ∑_*g*_ *x*_*cg*_)^2^] to the loss function. In models with ZINB, training stopped earlier than in models with ZIP. In terms of the MSE of the rates, the models with ZIP and ZINB were not significantly different (Fig. 11). However, the models with ZIP performed significantly better regarding NLL, and their first layer weights were much sparser (Fig. 12) than those with ZINB.

**Figure 11:**
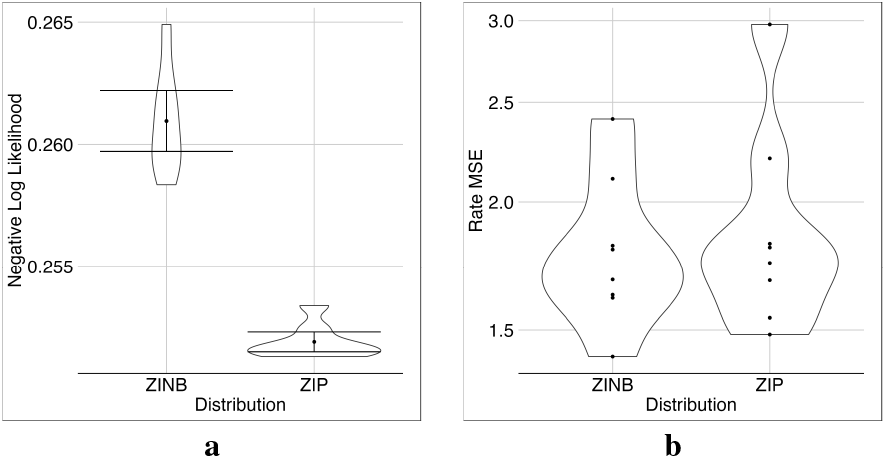
Comparing the performance of distributions ZINB vs ZIP. ZIP had better NLL than ZINB for the ROSMAP dataset **(a)** without sacrificing MSE **(b)**.

**Figure 12:**
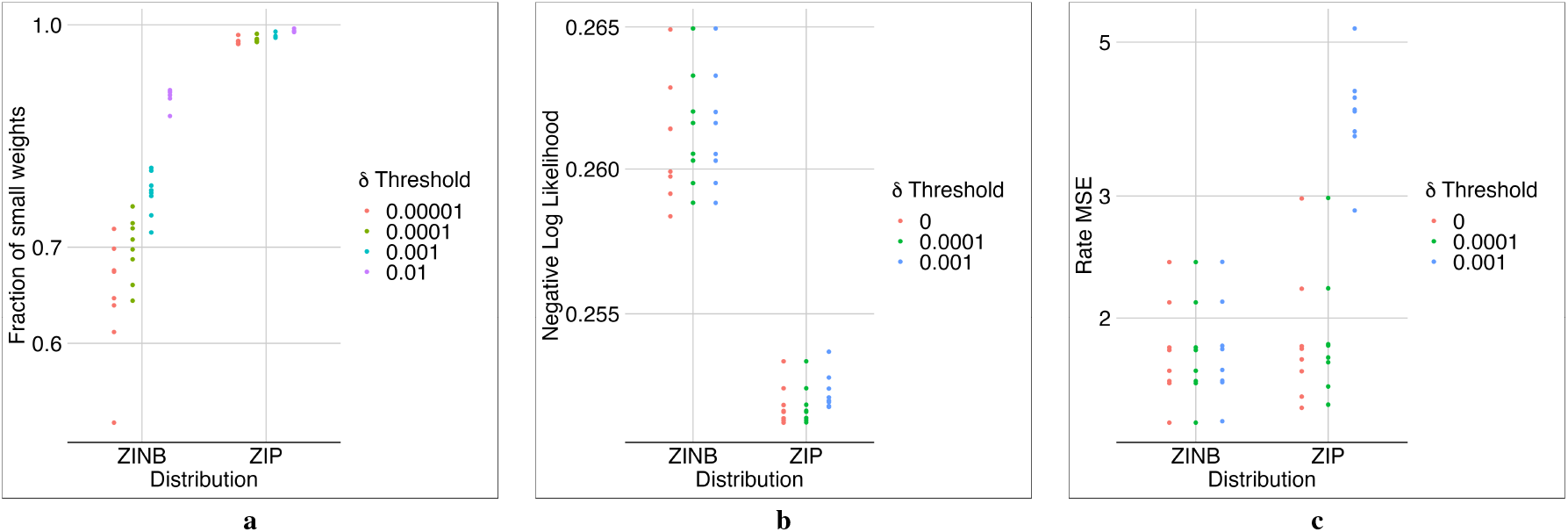
The effect of ZINB and ZIP distributions on the model’s sparsity. ZINB did not lead to sparsifiable weights **(a)**, choosing δ = 0.0001 for ZIP led to a very sparse model with almost the same performance before pruning (i.e., δ = 0.0001) **(b)**,**(c)**.

### G. ZiPo compared to DCA

Due to technical difficulties, we could not use the DCA package to compare the corresponding method with ZiPo. Instead, we leveraged our implementation, which is in a sense a generalization of DCA, to simulate DCA using three approaches: (1) the recommended parameters by DCA authors in our settings, (2) the Poisson distribution (zipDCA), and (3) the settings that are similar to our optimized ZiPo model (tunedDCA). For the base settings, we used a 128–64–128 structure for the number of neurons in the model, applied logarithmic transformation, and normalized the input matrix by dividing the expression of each gene by the library size of each cell.

In terms of NLL, ZiPo, and DCA performed similarly and better than other models (Fig. 13a). The simplest model, i.e., DCA, was trained faster than the others but had the largest MSE for the rates (Fig. 13b). Poisson distribution would decrease the rates’ MSE, but the NLL was not as low as ZiPo.

**Figure 13:**
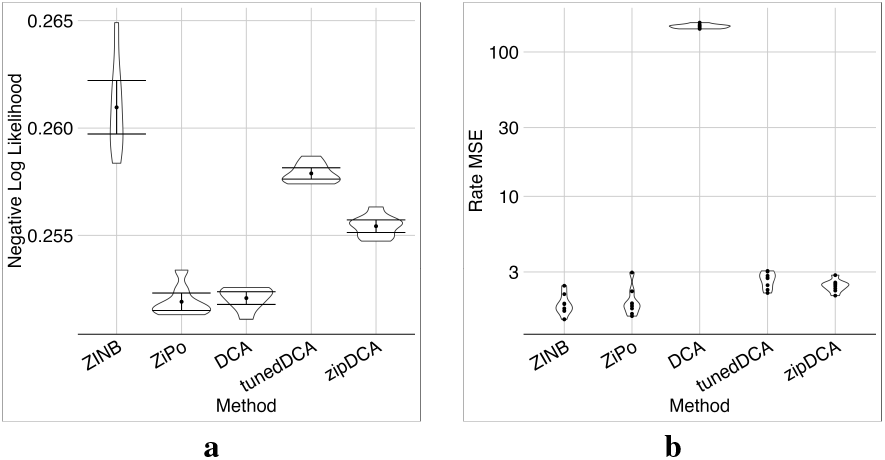
Comparing ZiPo and similar DCA methods. Negative log likelihood **(a)** and MSE **(b)** for different methods are shown. While pure DCA matched ZiPo in terms of NLL performance, its MSE results were notably inferior. Adjusting DCA parameters or opting for a Poisson distribution instead of a negative binomial distribution might improve the MSE, but this comes at the cost of diminishing NLL performance.

### H. Automatic hyperparameter optimization

We used the “Optuna” package to optimize ZiPo’s hyperparameters. Specifically, we searched five different encoder structures, 2048–1024–512–256–128, 1024–512–256–128, 1024– 512–256–128–64, 512–256–128, and 512–256–128–64 with full residual connections. For the decoder, we used a simple linear model For the regularization parameter of zero inflation and the weights, we searched the range 0.01–1 in the logarithmic space. After 96 iterations, the hyper-optimizer chose the deepest model, which resulted in a relatively reasonable value for the MSE of expression rates (Fig. 14). For the best model, the regularization parameter of zero inflation was 0.10594359 and the regularization parameter of weights was 0.14182903, both of which were near our choice of 0.1.

**Figure 14:**
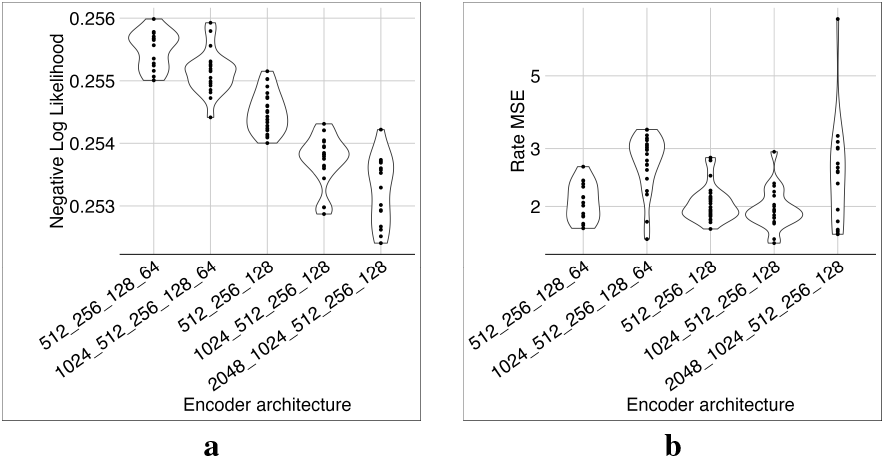
Hyperparameter search. The hyper-optimizer determined that the deepest model had the best NLL **(a)**, despite having similar MSE **(b)** as other models.

### I. Three additional datasets

Using the same hyperparameters optimized on the liver dataset, we trained the ZiPo model on the three datasets mentioned in II-F. The ZINB distribution generally seemed more powerful due to its larger number of parameters (Fig. 15). ZINB requires fewer epochs, which means faster convergence of the optimizer, most probably because it has more parameters and can easily fit the input distribution. However, ZIP acted even better regarding NLL in cases like the Liver dataset.

**Figure 15:**
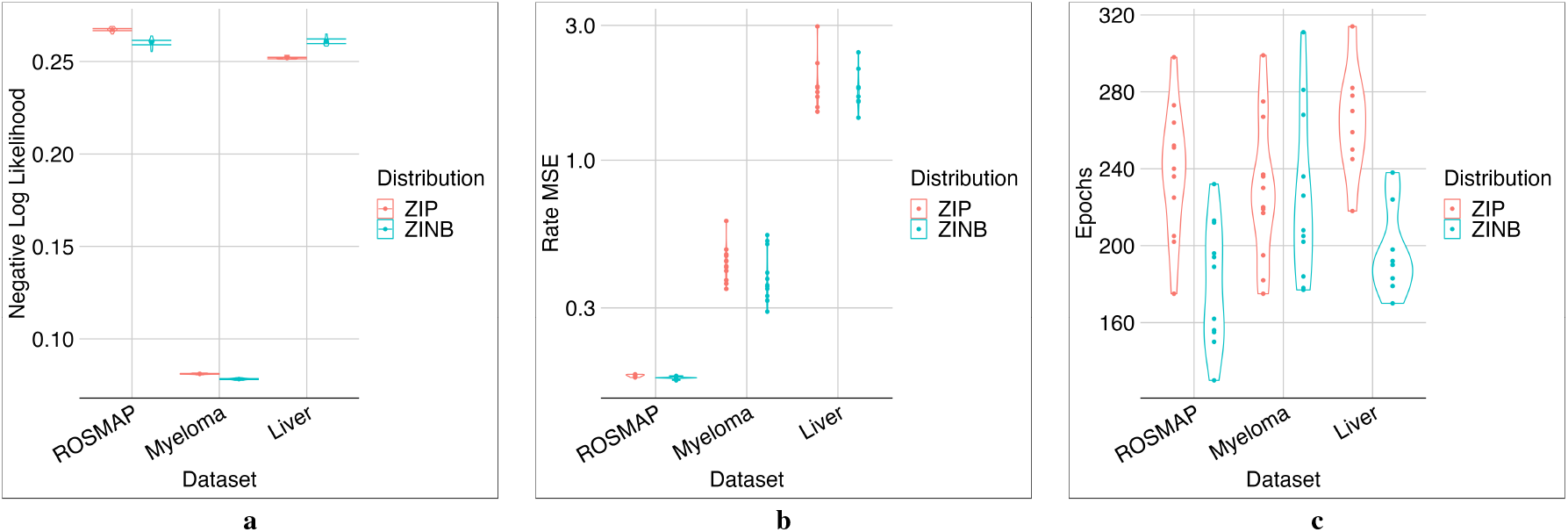
Comparing ZINB and ZIP in three datasets. The NLL **(a)** and MSE **(b)** of reconstructed rates are shown. For some datasets, ZIP might perform even better than ZINB. The model with ZINB would train faster, perhaps due to its richer parameter space **(c)**.

## IV. Discussion

We have described a novel deep model, ZiPo, for rate estimation and library size prediction in scRNA-seq data. ZiPo presents several unique features described below. Most prominently, the present model features an adjustable zeroinflation rate.

a. *Handling big data with deep models and batches*: A significant advantage of using deep models is their capability to handle massive datasets in batches. In contrast, other methods would be impractical as they would need to load the whole dataset in the memory. Also, similar to other deep models, ZiPo can be partially trained with one dataset and then fine-tuned using other datasets. This usually cannot be done easily with other methods.
b. *Software quality*: To ensure the longevity of our code and its maintenance in the long term, we have used popular software packages and modern programming techniques in implementing ZiPo. Using features like configuration files, we have provided a flexible platform for software developers and end-users.
c. *ZIP overall surpasses ZINB*: With an appropriate architecture, zero-inflation models using ZIP distribution perform well. As shown here, the models with ZINB distribution can achieve smaller losses, hence better performances than those using ZIP. However, due to the need for extra regularization in the models with ZINB, they are relatively larger with more trainable weights, their training is more complicated and time-consuming, and they are harder to interpret than those with ZIP.
d. *Latents for downstream analysis*: To assess the validity of the latent variables, we tested ZiPo for clustering the cells in the ROSMAP dataset. We compared the results with the predefined cell types. The tSNE plots [39] based on the latent variables from the two ZIP and ZINB models showed that these models generally performed well except for a few outliers (Fig. 16).

**Figure 16:**
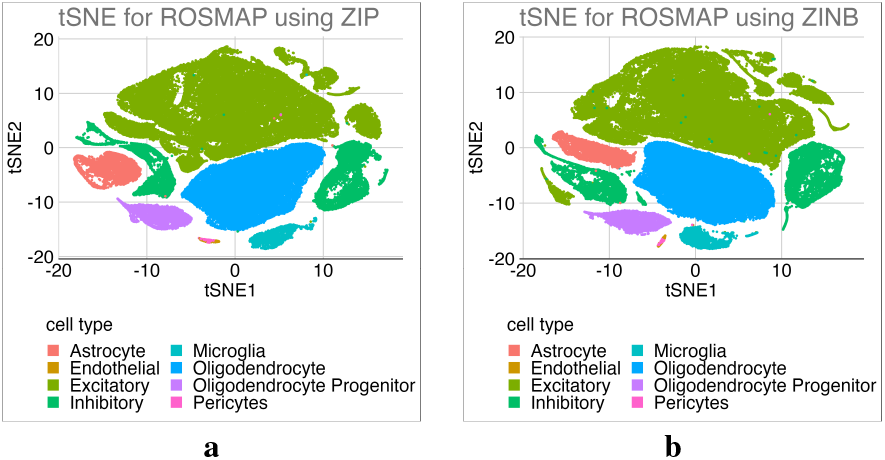
tSNE plots for latent variables. tSNE plots for the resulting latent variables of ZiPo using ZIP **(a)** and ZINB **(b)** distributions are shown. Cell types were previously determined in the ROSMAP dataset.
e. *Future work*: The latent variables can be investigated more thoroughly and used in downstream analyses including regressing eigengenes [40] or classifying senescent cells [41].

## Acknowledgments

The ROSMAP data were provided by the Rush Alzheimer’s Disease Center, Rush University Medical Center, Chicago. Generation of this dataset was supported by the National Institute on Aging (NIA) grants RF1AG57473, P30AG010161, R01AG015819, R01AG017917, U01AG46152, U01AG61356, RF1AG059082, P30AG072975, and R01AG036042. Additional phenotypic ROSMAP data can be requested at https://www.radc.rush.edu. We acknowledge the Texas Advanced Computing Center (TACC) at The University of Texas at Austin for providing high-performance computing (HPC) resources (http://www.tacc.utexas.edu).

## Author Contributions

M.S. contributed most of the ideas, developed the code, and prepared the experiments. S.K. conducted experiments. J.R. helped with figure preparation and coding. H.Z and M.S. analyzed the results. M.S., S.K., S.S., and H.Z. wrote the manuscript. S.S. contributed to the conduction of the experiments and writing the manuscript. All authors reviewed the manuscript.

## Conflict of Interest Disclosures

The authors have declared that no competing interests exist.

---

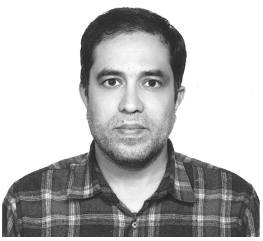

**Mohsen Sharifitabar** is a mathematician with a wealth of experience in optimization, computer programming, and big data analysis. He received his PhD from the Department of Mathematical Sciences at Sharif University of Technology, Iran. His current work is focused on developing machine learning techniques for analyzing large biological datasets, including genome-wide association studies, multiomics, and analysis of single-cell and spatial omics. He is now contributing to various projects requiring the analysis of different types of biological data.

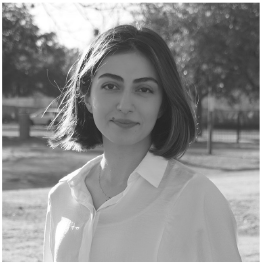

**Shiva Kazempour** is a PhD candidate at The University of Texas Health Science Center at San Antonio. Her research interests include discovering the factors contributing to the brain aging. Her current research is focused on developing deep learning models for the identification of senescent cells in neurodegenerative diseases using spatial transcriptomics and proteomics.

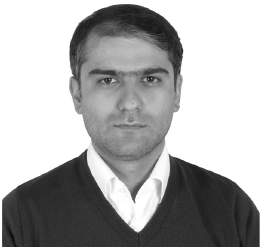

**Javad Razavian** earned his M.Sc. in Computer Science from Sharif University of Technology in 2007. He currently serves as a Lecturer in the Department of Computer Science at the University of Qom and as a Volunteer Researcher at Oncinfo Research Lab, UT Health San Antonio, Texas. His research focuses on developing advanced machine learning models and exploring applications of deep learning that directly impact human life.

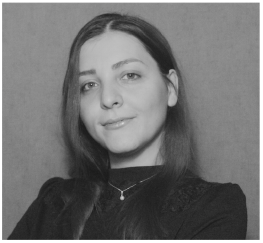

**Sogand Sajedi** received her Ph.D. from the Vienna University of Technology, Austria. She is currently a research scientist at MD Anderson Cancer Center, Texas. Her research is centered on multi-omics analysis in cancer, emphasizing integrating genomic and transcriptomic single-cell datasets.

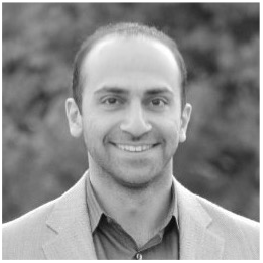

**Soroosh Solhjoo** is an expert in processing biological signals, computational modeling, and developing machine-learning techniques for data analysis. His research interests include bioenergetics, cardiovascular health, and creating tools for single-cell RNA sequencing analysis. He received his PhD in Biomedical Engineering from the Johns Hopkins University School of Medicine in Baltimore, Maryland.

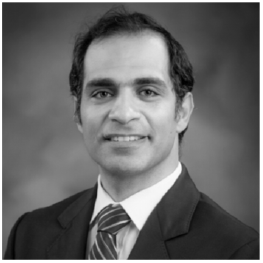

**Habil Zare** is a computational biologist. He received his PhD from the Computer Science Department at the University of British Columbia in Canada. His current research is focused on applications of machine learning in mining large biological datasets, including gene network analysis, integrating multiomics, and analysis of single-cell and spatial omics. Through inferring important biological information from these datasets, his lab has contributed to several studies on dementia and cancer research.

## Notes

This work was supported in part by NIH-NIA under Grants R01AG057896, 1RF1AG063507, R01AG068293, 1R01AG0665241A, 1R01AG065301, and P30AG066546, and in part by NIH-NINDS under Grants RF1NS112391 and U19NS115388.

https://bitbucket.org/habilzare/alzheimer/src/master/code/deep/ZiPo

### Competing Interest Statement

The authors have declared no competing interest.

